# Emergence of NADP^+^-reducing enzymes in *Escherichia coli* central metabolism via adaptive evolution

**DOI:** 10.1101/2021.02.08.430375

**Authors:** Madeleine Bouzon, Volker Döring, Ivan Dubois, Anne Berger, Gabriele M. M. Stoffel, Liliana Calzadiaz Ramirez, Sophia Meyer, Marion Fouré, David Roche, Alain Perret, Tobias J. Erb, Arren Bar-Even, Steffen N. Lindner

## Abstract

The nicotinamide cofactor specificity of enzymes plays a key role in regulating metabolic processes and attaining cellular homeostasis. Multiple studies have used enzyme engineering tools or a directed evolution approach to switch the cofactor preference of specific oxidoreductases. However, whole-cell adaptation towards the emergence of novel cofactor regeneration routes was not previously explored. To address this challenge, we used an *Escherichia coli* NADPH-auxotroph strain. We continuously cultivated this strain under selective conditions. After 500-1100 generations of adaptive evolution using different carbon sources, we isolated several strains capable of growing without an external NADPH source. Most isolated strains were found to harbor a mutated NAD-dependent malic enzyme (MaeA). A single mutation in MaeA was found to switch cofactor specificity while lowering enzyme activity. Most mutated MaeA variants also harbored a second mutation that restored the catalytic efficiency of the enzyme. Remarkably, the best MaeA variants identified this way displayed overall superior kinetics relative to the wildtype variant with NAD^+^. In other evolved strains, the dihydrolipoamide dehydrogenase (Lpd) was mutated to accept NADP^+^ thus enabling the pyruvate dehydrogenase and 2-ketoglutarate dehydrogenase complexes to regenerate NADPH. Interestingly, no other central metabolism oxidoreductase seems to evolve towards reducing NADP^+^, which we attribute to several biochemical constraints such as unfavorable thermodynamics. This study demonstrates the potential and biochemical limits of evolving oxidoreductases within the cellular context towards changing cofactor specificity, further showing that long-term adaptive evolution can optimize enzyme activity beyond what is achievable via rational design or directed evolution using small libraries.

**Importance:** In the cell, NAD(H) and NADP(H) cofactors have different functions. The former mainly accepts electrons from catabolic reactions and carries them to respiration, while the latter provides reducing power for anabolism. Correspondingly, the ratio of the reduced to the oxidized form differs for NAD (low) and NADP (high), reflecting their distinct roles. We challenged the flexibility of *E. coli’s* central metabolism in multiple adaptive evolution experiments using an NADPH-auxotroph strain. We found several mutations in two enzymes, changing the cofactor preference of malic enzyme and dihydrolipoamide dehydrogenase. Upon deletion of their corresponding genes we performed additional evolution experiments which did not lead to the emergence of any additional mutants. We attribute this restricted number of mutational targets to intrinsic thermodynamic barriers: The high ratio of NADPH to NADP^+^ limits metabolic redox reactions which can regenerate NADPH, mainly by mass action constraints.

## Introduction

The cofactor preference of enzymes is crucial for ensuring balanced production and consumption of resources, proper regulation of metabolic processes, and general cellular homeostasis. The two primary electron carriers, NAD and NADP, demonstrate this quite well, as the former is mainly involved in catabolic and respiratory processes while the latter mostly participates in biosynthetic pathways. The physiological reduction levels of NAD and NADP pools reflect this distinction: the NAD pool is highly oxidized, providing a thermodynamic push for catabolic processes, which are mostly oxidative and use NAD^+^ as an electron acceptor; in contrast, the NADP pool is relatively reduced, thermodynamically supporting anabolic processes, which are mostly reductive, using NADPH as an electron donor.

Multiple previous studies have demonstrated how the replacement of only a few residues in the active site of an oxidoreductase enzyme can switch its cofactor preference from NAD to NADP or *vice versa* [1-4]. Several studies also constructed mutant libraries of specific oxidoreductases and harnessed natural selection to identify variants with altered cofactor specificity [5-7]. Yet, until now, no study has attempted to systematically explore the overall evolvability of central metabolism oxidoreductases towards the use of a different cofactor. This could help shed light on the flexibility of the metabolic network and identify emergent regeneration processes, thus adding to our understanding on the plasticity of cellular metabolism.

Here, we applied a whole cell evolution approach towards the emergence of new NADPH regeneration routes. We used an NADPH-auxotroph strain, deleted in all enzymes capable of regenerating NADPH (Δ*zwf* Δ*maeB* Δ*icd* Δ*pntAB* Δ*sthA*), with the exception of 6-phosphogluconate dehydrogenase [8]. This strain could grow on a minimal medium only when gluconate is added as an NADPH source. We conducted multiple parallel evolution experiments in continuous culture with limiting supply of gluconate, thus selecting for the emergence of mutated oxidoreductases that could reduce NADP^+^. After long cultivation periods under selective conditions (500-1100 generations), we were able to isolate evolved strains, from 10 independent evolution experiments, capable of growing without gluconate. Despite conducting the experiments with different carbon sources, each enforcing a different distribution of central metabolism fluxes, we found that all the evolved strains harbor mutations in one of only two enzymes: NAD-dependent malic enzyme (MaeA) or dihydrolipoamide dehydrogenase (Lpd). We show that while the mutated MaeAs strongly prefer the reduction of NADP^+^ over NAD^+^, the catalytic efficiencies of the Lpd variants are in similar ranges with both cofactors. Several mutated MaeA variants also displayed higher overall activity than their wildtype counterpart, demonstrating the strength of selective adaptation for identifying superior mutants. Interestingly, no other central metabolism oxidoreductase evolved towards the use of NADP^+^, which could be attributed to structural or thermodynamic constraints.

## Results

### The NADPH-auxotroph strain and oxidoreductase candidates for NADPH regeneration

Five enzymes are known to support NADPH regeneration in *E. coli*: glucose 6-phosphate dehydrogenase (Zwf), 6-phosphogluconate dehydrogenase (Gnd), the NADP-dependent malic enzyme (MaeB), isocitrate dehydrogenase (Icd), and the membrane-bound transhydrogenase (PntAB) [9]. In a previous study we constructed a strain in which the genes coding for these NADP-dependent oxidoreductases were deleted, with the exception of *gnd* (Δ*zwf* Δ*maeB* Δ*icd* Δ*pntAB* Δ*sthA*; the gene *sthA*, which encodes the soluble transhydrogenase, was also deleted to remove a major NADPH sink) [8]. For this NADPH-auxotroph strain to grow on a minimal medium, gluconate must be added as a precursor of 6-phosphogluconate, the substrate of Gnd. Due to the deletion of *icd*, the supply of 2-ketoglutarate as a precursor for glutamate and the downstream C_5_ amino acids glutamine, proline, and arginine is also mandatory. We demonstrated that, when gluconate is omitted, the NADPH-auxotroph strain can serve as an effective *in vivo* platform to test and optimize different enzymatic systems for NADPH regeneration [8].

We speculated that cultivating the NADPH-auxotroph strain under limiting amounts of gluconate would lead to the emergence of mutated oxidoreductase enzymes capable of regenerating NADPH. Such an enzyme would need to sustain very high flux to support a NADPH regeneration rate sufficiently high to enable cell growth. We therefore regarded oxidoreductase enzymes that participate in central metabolism as the major candidates for evolution towards NADP^+^ reduction. Central metabolism employs multiple oxidoreductases (Fig. 1), including glyceraldehyde 3-phosphate dehydrogenase (GapA), glycerol dehydrogenase (GldA), glycerol 3-phosphate dehydrogenase (GpsA), pyruvate/2-ketoglutarate dehydrogenases (or, more precisely, their lipoamide dehydrogenase subunit, Lpd), lactate dehydrogenase (LdhA), the NAD-dependent malic enzyme (MaeA), and malate dehydrogenase (Mdh).

**Figure 1:**
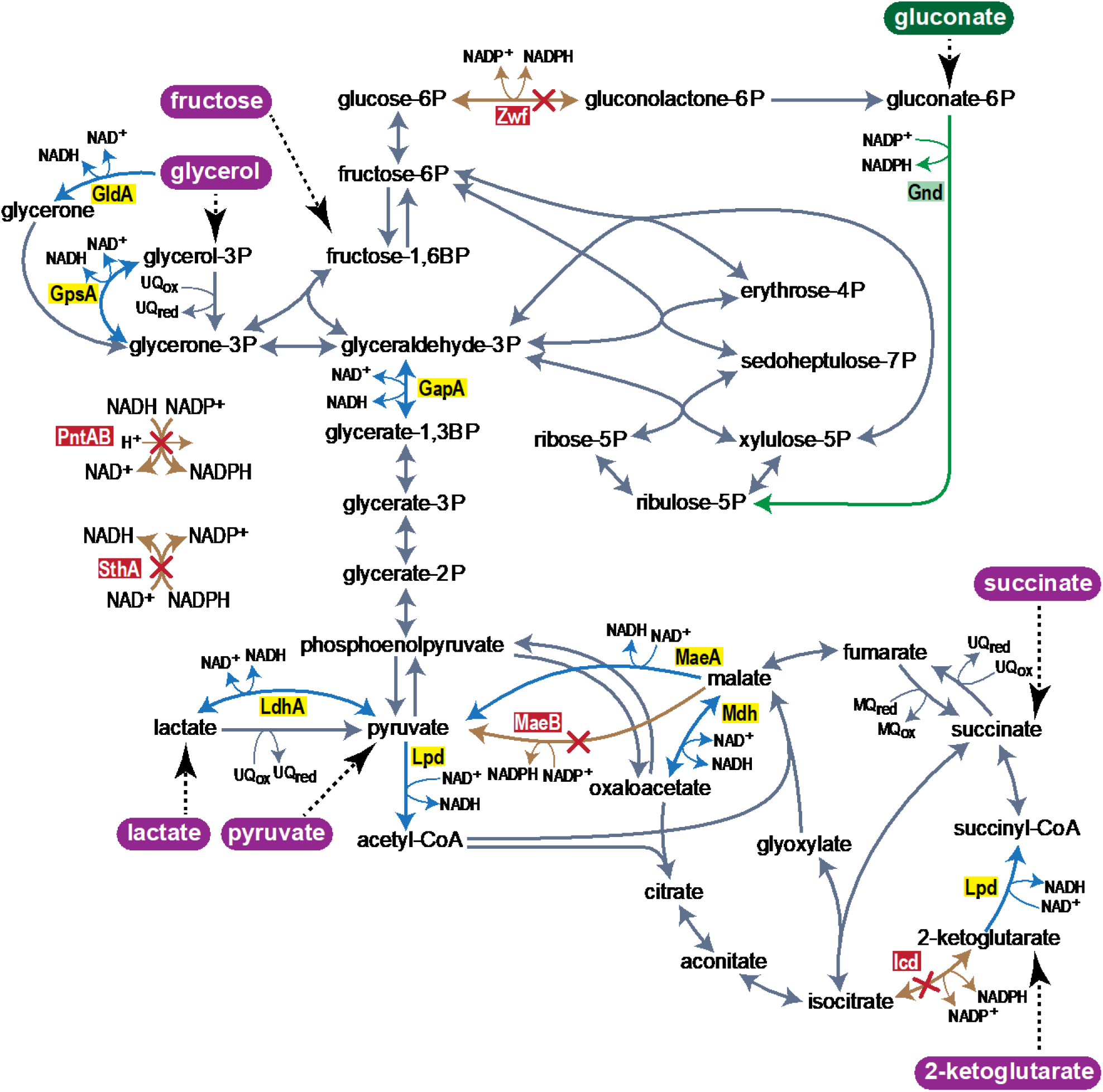
Central carbon metabolism of *E. coli*. NADP^+^ reducing reactions, which were deleted to construct the NADPH-auxotroph strain are shown by red crossed orange arrows and red underlain enzyme name. The green arrow indicates gluconate dependent NADPH generation. Blue arrows combined with yellow underlain enzyme names highlight NAD^+^-dependent oxidoreductases and potential candidates for a cofactor change from NAD^+^ to NADP^+^ in evolution experiments. Highlighted in purple are carbon sources used in evolution experiments. Abbreviations of enzymes: Zwf, glucose 6-phosphate dehydrogenase; Gnd, gluconate 6-phosphate dehydrogenase; GldA, glycerol dehydrogenase; GpsA, glycerol 3-phosphate dehydrogenase; GapA, glyceraldehyde 3-phosphate dehydrogenase; PntAB, membrane bound transhydrogenase; SthA, soluble transhydrogenase; LdhA, lactate dehydrogenase; Lpd, dihydrolipoamide dehydrogenase; MaeB, NADP^+^-dependent malic enzyme; MaeA, NAD^+^-dependent malic enzyme; Mdh, malate dehydrogenase; Icd, isocitrate dehydrogenase.

Since the entry point of carbon into central metabolism dictates the carbon flux distribution, we hypothesized that the choice of the carbon source could predispose different NAD^-^dependent oxidoreductases as targets for mutations. For example, mutations in GapA that enable it to accept NADP^+^ would be useful to regenerate NADPH only if the cell is fed with a carbon source that enters upper glycolysis and induces glycolytic flux (rather than gluconeogenesis). Similarly, mutagenesis of GpsA or LdhA towards accepting NADP^+^ could effectively produce NADPH only when glycerol or lactate (respectively) serves as the carbon source.

### Adaptive evolution of the NADPH-auxotroph strain led to mutations in *maeA* and *lpd*

We conducted twelve evolution experiments using six carbon sources: fructose, glycerol, pyruvate, lactate, 2-ketoglutarate and succinate (two parallel cultures for each carbon source). Fructose and glycerol are expected to force glycolytic and anaplerotic fluxes, pyruvate and lactate are expected to force gluconeogenic and anaplerotic fluxes, while 2-ketoglutarate and succinate are expected to force gluconeogenic and cataplerotic fluxes (cataplerosis being the reverse of anaplerosis, that is, decarboxylation of a C_4_ intermediate of the TCA cycle to generate a C_3_ glycolytic intermediate). Hence, the six carbon sources nicely cover a large variation in flux distribution across central metabolism (Fig. 1).

We used GM3 cultivation devices to apply a medium-swap continuous culture regime [10, 11] in order to evolve the NADPH-auxotroph strain towards novel NADPH regeneration routes. Cultures of growing cells subjected to this regime are diluted, at fixed time intervals, by one of two growth media, permissive or stressing, the choice depending on the turbidity of the culture measured in real time. Specifically, if the turbidity is below a predefined value, a dilution pulse comes from the permissive medium; otherwise, the stressing medium is used to dilute the culture [10, 11]. This approach enables gradual genetic adaptation of a bacterial population to grow on the stressing medium. Here, the permissive medium contained one of the six canonical carbon sources listed above, gluconate as NADPH source, and 2-ketoglutarate as glutamate source. The stressing medium had the same composition except for gluconate which was omitted. Continuous cultivation under these conditions is expected to select for the emergence of novel NADPH regenerating enzymes, adapting the cells to grow with less and less gluconate, until growth on the stressing medium alone is reached.

Of the 12 parallel adaptive evolution experiments, 8 evolved to rely completely on the stressing medium (100% stressing medium pulses), including at least one culture for all six carbon sources used (Fig. 2A). The adaptation kinetics and the number of generations required to attain growth without gluconate were comparable for all the eight cultures (Table 1). In most cases, the stressing/relaxing dilution ratio only slightly increased during a prolonged period of the adaptation until a sharp rise occurred and growth on the stressing medium was attained, pointing to the appearance of a key adaptive mutation in the population.

**Table 1.**
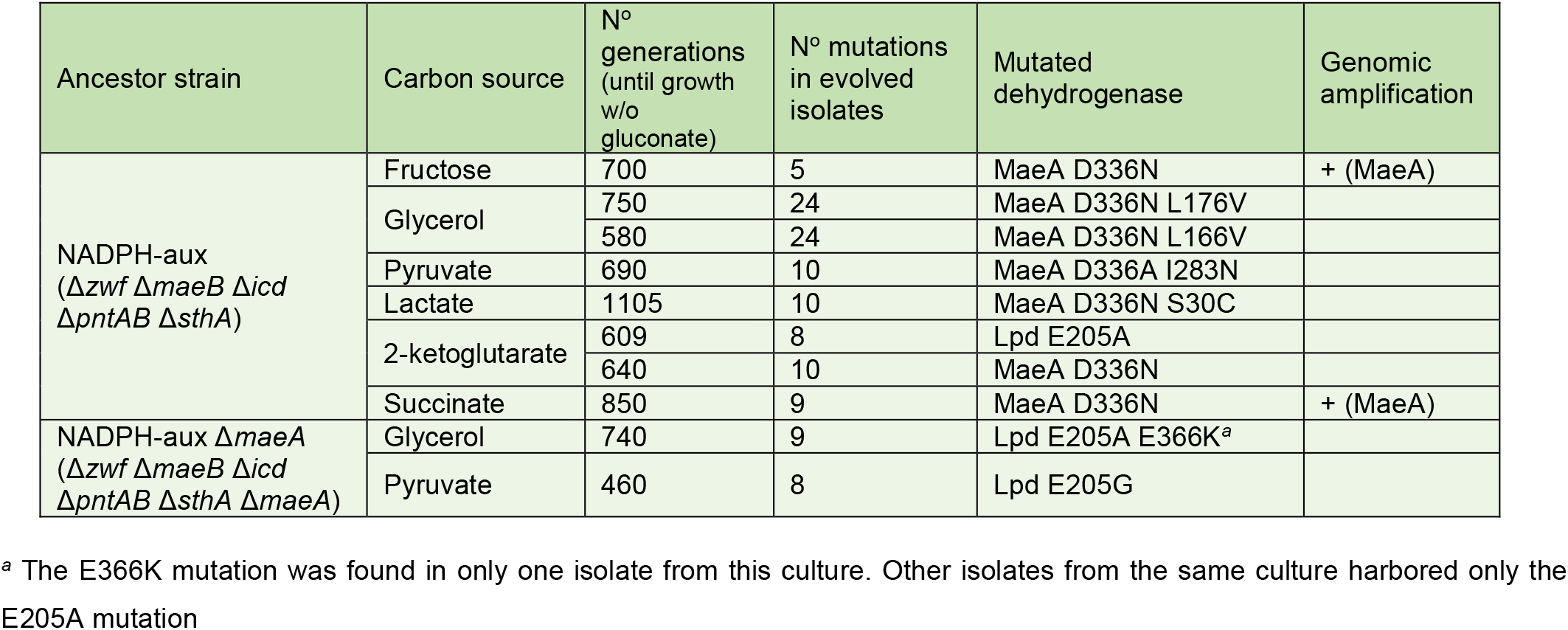
Outcome of the successful adaptive evolutions of NADPH-auxotroph strains performed in the GM3 cultivation device. See Supplementary Data, for all identified mutations.

| Ancestor strain | Carbon source | N° generations (until growth w/o gluconate) | N° mutations in evolved isolates | Mutated dehydrogenase | Genomic amplification |
| --- | --- | --- | --- | --- | --- |
| NADPH-aux<br>($\Delta zwf \Delta maeB \Delta icd \Delta pntAB \Delta sthA$) | Fructose | 700 | 5 | MaeA D336N | + (MaeA) |
|  | Glycerol | 750 | 24 | MaeA D336N L176V |  |
|  |  | 580 | 24 | MaeA D336N L166V |  |
|  | Pyruvate | 690 | 10 | MaeA D336A I283N |  |
|  | Lactate | 1105 | 10 | MaeA D336N S30C |  |
|  | 2-ketoglutarate | 609 | 8 | Lpd E205A |  |
|  |  | 640 | 10 | MaeA D336N |  |
|  | Succinate | 850 | 9 | MaeA D336N | + (MaeA) |
| NADPH-aux $\Delta maeA$<br>($\Delta zwf \Delta maeB \Delta icd \Delta pntAB \Delta sthA \Delta maeA$) | Glycerol | 740 | 9 | Lpd E205A E366K<sup>a</sup> | |
|  | Pyruvate | 460 | 8 | Lpd E205G |  |

<sup>a</sup> The E366K mutation was found in only one isolate from this culture. Other isolates from the same culture harbored only the E205A mutation

**Figure 2:**
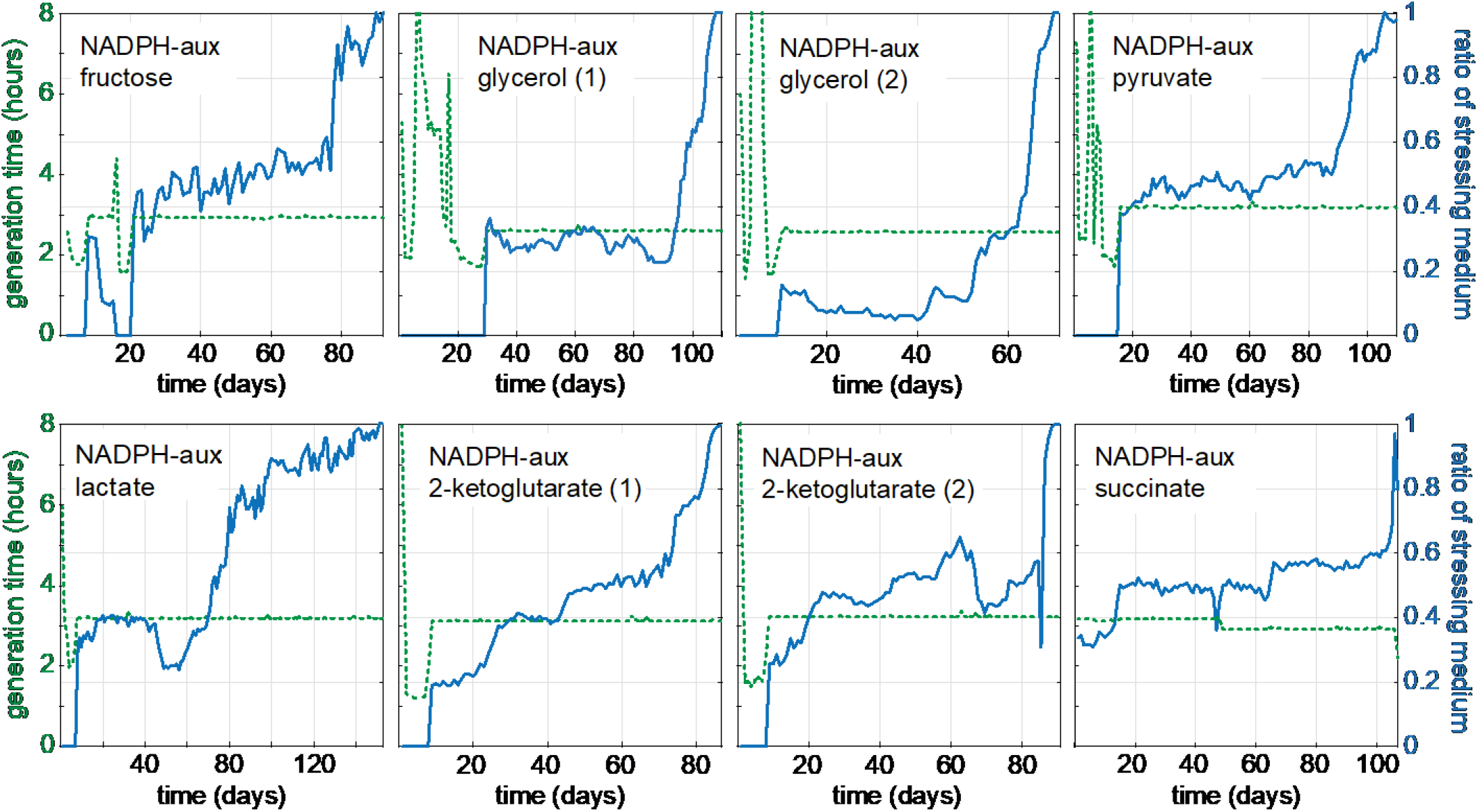
Evolution of the NADPH-auxotroph strain for growth with either fructose, glycerol, pyruvate, lactate, 2-ketoglutarate, or succinate in the absence of gluconate. For each carbon source 2 independent cultures were subjected to a medium swap regime in GM3 devices (see methods section). Shown are the eight cultures which evolved to growth in stressing medium. Blue lines show the ratio of stressing medium over relaxing medium. Stressing media contained 5 mM 2-ketoglutarate plus one of the following carbon sources: D-fructose (10mM), succinate (17 mM), pyruvate (25 mM), lactate (25 mM), glycerol (20mM), 2-ketoglutarate (20mM final)) (right axes). Relaxing media were composed as stressing medium supplemented with 5 mM D-gluconate. Generation times are indicated by the green dashed lines (left axes).

We isolated strains from all the cultures that were evolved to grow on the stressing medium and cultivated each of these isolated strains on a minimal medium supplemented with different carbon sources (Fig. 3). Each of these strains could grow well on the carbon source used in the evolution experiment in which it has emerged. The isolated strains could also grow on (almost) all other carbon sources tested, indicating that the metabolic adaptation was not restricted to a particular flux distribution in central metabolism.

**Figure 3:**
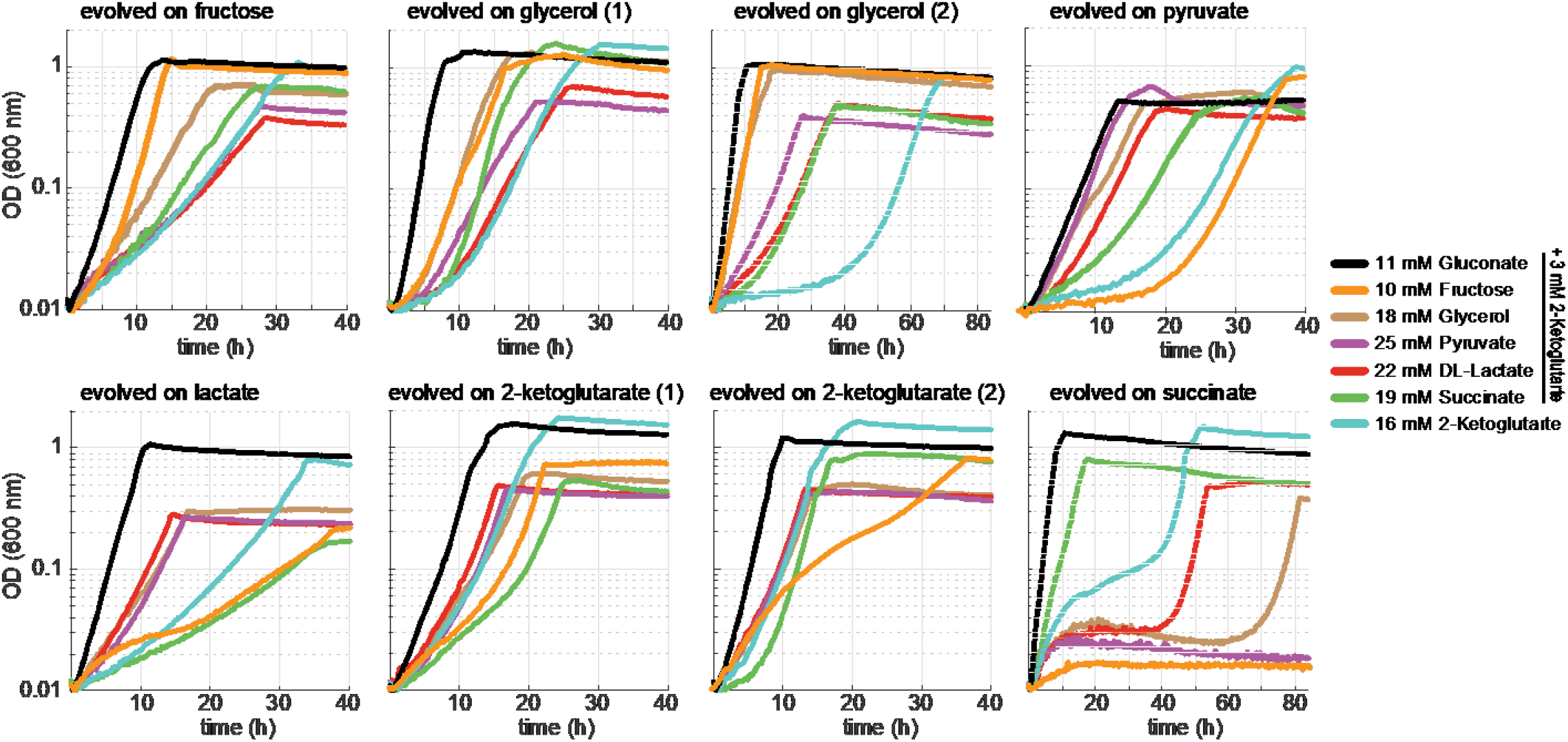
Growth curves of isolates obtained from cultures adapted in GM3 to proliferate without gluconate on either fructose, glycerol, pyruvate, lactate, 2-ketoglutarate, or succinate, each supplemented with 5 mM 2-ketoglutarate. Growth of the isolates was determined on 11 mM gluconate, 10 mM fructose, 18 mM glycerol, 25 mM pyruvate, 22 mM D/L-lactate, 19 mM succinate (all supplemented with 3 mM 2-ketoglutarate), and 16 mM 2-ketoglutarate. Growth curves were recorded in triplicates, showing similar growth (± 5%).

Genomic sequencing of the adapted strains and comparison with the non-evolved parent strain revealed 5 to 10 point mutations as well as small insertions and deletions in all genomes sequenced (Table 1). Importantly, we sequenced two isolates from each successful adaptive evolution experiment; for each experiment, these isolates displayed an almost identical mutation profile (Supplementary Data), suggesting that the bacterial populations in the cultures were rather homogeneous. The strains isolated from the two glycerol cultures were exceptional as each contained more than 20 mutations, including a missense mutation in gene *mutL* coding for a DNA mismatch repair protein. Notably, in 7 of the 8 cultures, the gene *maeA*, which codes for the NAD-dependent malic enzyme, carried one or two non-silent mutations. Furthermore, the isolates from the fructose and succinate cultures carried an amplified chromosomal region containing the *maeA* gene, which points to overexpression of the mutated gene as an additional adaptive trait. The only divergent isolate was from one of the cultures cultivated on 2-ketoglutarate, in which *maeA* was not mutated. Instead, *lpd*, coding for lipoamide dehydrogenase, was mutated in this strain.

### A mutation in a single residue in MaeA changed cofactor specificity but other mutations were essential to recover catalytic efficiency

Three of the isolated strains harbored a single mutation in *maeA*: D336N. In four strains, *maeA* had two mutations, of which one was either D336N or D336A (Table 1). We chose to focus on three mutated variants: D336N, D336N L176V, and D336A I283N. We introduced each of these mutation sets into the non-evolved, parental strain using Multiplex Automated Genomic Engineering (MAGE [12]) and characterized the growth of the resulting strains (Fig. 4).

**Figure 4:**
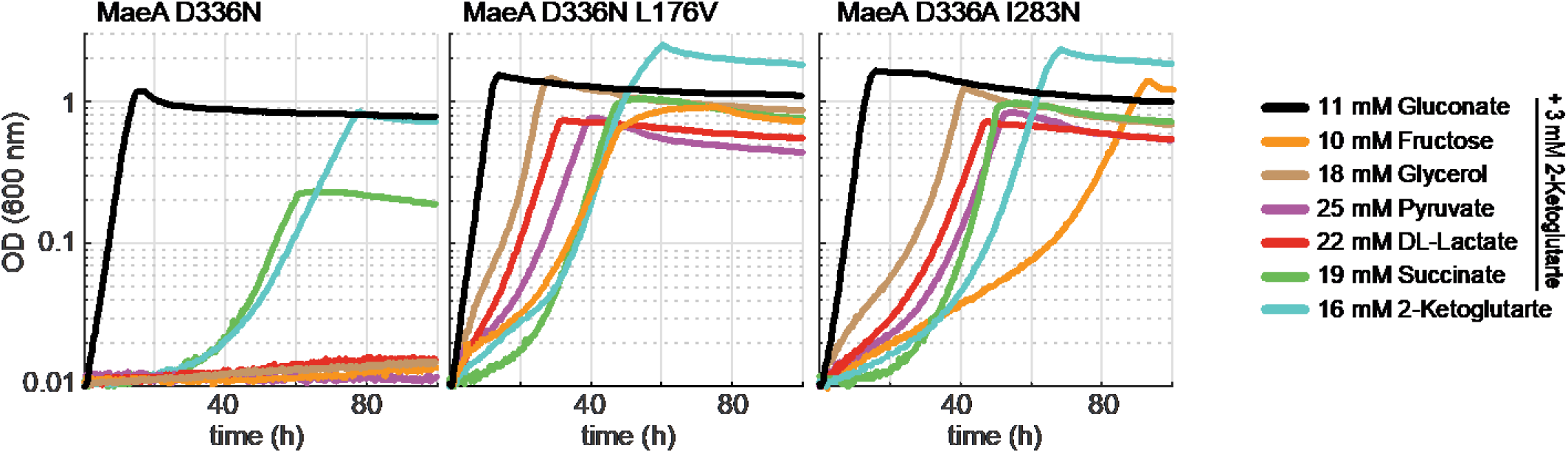
Growth of NADPH-auxotroph strain derivatives carrying mutations in *maeA* (D336N, D336N + L176V, D336A + I283N) introduced by MAGE. Growth was determined on 11 mM gluconate, 10 mM fructose, 18 mM glycerol, 25 mM pyruvate, 22 mM D/L-lactate, 19 mM succinate (all supplemented with 3 mM 2-ketoglutarate), and 16 mM 2-ketoglutarate. Growth curves were recorded in triplicates, showing similar growth (± 5%).

The strain harboring *maeA* D336N was able to grow only with succinate and 2-ketoglutarate but not with the other carbon sources (Fig. 4A). This is in line with the fact that two of the three evolved strains displaying this mutation were cultivated on either succinate or 2-ketoglutarate, while the third one – cultivated on fructose – also showed an amplification of the chromosomal region containing the *maeA* gene (Table 1). It therefore seems that the D336N mutation enhanced NADP^+^ reduction by MaeA, but only to a limited extent. Therefore, only carbon sources that enter the TCA cycle (i.e., succinate and 2-ketoglutarate) and thus force high cataplerotic flux via MaeA, support sufficiently high NADPH regeneration rate. When another carbon source is used (e.g., fructose) overexpression of *maeA* D336N seems necessary to enable sufficient NADPH regeneration.

On the other hand, the strains harboring either *maeA* D336N L176V or *maeA* D336A I283N were able to grow on (almost) all carbon sources (Fig. 4B,C). This suggests that these mutation sets increased the activity of MaeA with NADP^+^ to a sufficiently high level such that even carbon sources that do not induce cataplerotic flux could sustain high NADPH regeneration rate without further overexpression of *maeA*.

To test whether these interpretations are correct, we purified the mutated MaeA variants and performed steady state analysis with NAD^+^ and NADP^+^ (Table 2). We found that while wildtype (WT) MaeA can accept NADP^+^, it uses this cofactor with a low *k*_*cat*_/*K*_*M*_ = 11 s^-1^ mM^-1^, more than two orders of magnitude lower than the *k*_*cat*_/*K*_*M*_ > 1800 s^-1^ mM^-1^ measured with NAD^+^. The D336N mutation lowered *k*_*cat*_/*K*_*M*_ with NAD^+^ to 49 s^- 1^ mM^-1^, while increasing *k*_*cat*_/*K*_*M*_ with NADP^+^ ≈80-fold to ≈870 s^-1^ mM^-1^. This mutation thus increased MaeA preference toward NADP – as indicated by the ratio (*k*_*cat*_/*K*_*M*_)_NADP_/(*k*_*cat*_/*K*_*M*_)_NAD_ – by a factor of 3,000, from 0.006 to ≈18.

**Table 2.** Apparent steady state parameters of MaeA variants. Parameters are indicated as mean value ± standard error. Underlying Michaelis-Menten kinetics can be found in Supplementary Figure S1.

| Enzyme | Substrate | $K_M$ (mM) | $k_{cat}$ (s<sup>-1</sup>) | $k_{cat} / K_M$<br>(s<sup>-1</sup> mM<sup>-1</sup>) | $(k_{cat} / K_M)_{NADP} /$<br>$(k_{cat} / K_M)_{NAD}$ |
| --- | --- | --- | --- | --- | --- |
| MaeA WT | L-malate | 0.63 ± 0.04 |  |  | 0.0059 |
|  | NAD<sup>+</sup> | 0.10 ± 0.02 | 188 ± 9 | 1843 |  |
|  | NADP<sup>+</sup> | 3.6 ± 0.4 | 39 ± 2 | 11 |  |
| MaeA D336N | L-malate | 1.8 ± 0.2 |  |  | 17.6 |
|  | NAD<sup>+</sup> | 1.5 ± 0.2 | 74 ± 3 | 49 |  |
|  | NADP<sup>+</sup> | 0.091 ± 0.01 | 79 ± 2 | 868 |  |
| MaeA D336N L176V | L-malate | 1.4 ± 0.3 |  |  | 16.7 |
|  | NAD<sup>+</sup> | 0.27 ± 0.03 | 105 ± 2 | 389 |  |
|  | NADP<sup>+</sup> | 0.016 ± 0.001 | 102 ± 1 | 6497 |  |
| MaeA D336A I283N | L-malate | 0.48 ± 0.04 |  |  | 18.0 |
|  | NAD<sup>+</sup> | 0.27 ± 0.03 | 130 ± 3 | 480 |  |
|  | NADP<sup>+</sup> | 0.013 ± 0.00 | 112 ± 2 | 8615 |  |

The combined D336N L176V and D336A I283N mutations increased the activity with NADP^+^ even more, resulting in *k*_*cat*_/*K*_*M*_ of ≈6500 s^-1^ mM^-1^ and ≈8600 s^-1^ mM^-1^ (respectively), a 600- to 800-fold increase relative to MaeA WT. Interestingly, for these two mutation sets, the catalytic efficiency with NAD^+^ increased to the same extent as with NADP^+^, relative to that observed in MaeA D336N. Hence, the preference of MaeA D336N L176V and MaeA D336A I283N toward NADP^+^ was effectively identical to that of MaeA D336N. It therefore seems that the main role of the L176V and I283N mutations is the recovery of the catalytic efficiency lost upon cofactor switching by the mutation of D336 [1, 13]. Notably, the *k*_*cat*_/*K*_*M*_ values of MaeA D336N L176V and MaeA D336A I283N with NADP^+^ are ≈4-fold higher than the *k*_*cat*_/*K*_*M*_ of MaeA WT with NAD^+^, presenting one of rare cases in which overall relative catalytic efficiency was improved upon switching the cofactor specificity.

### Upon deletion of *maeA*, adaptive evolution led to mutations in *lpd*

As 7 of the 8 evolved strains contained a mutation in *maeA*, we decided to delete this gene in the NADPH-auxotroph strain and repeat the evolution experiment in the hope to prompt the emergence of other mutations enabling NADPH regeneration. Four cultures of the Δ*zwf* Δ*maeB* Δ*icd* Δ*pntAB* Δ*sthA* Δ*maeA* strain, two supplemented with pyruvate and two with glycerol, were cultivated under the medium swap continuous culture regime described above. For both carbon sources, growth on gluconate-free stressing medium was attained for one of the two parallel cultures (Fig. 5A). The culture supplemented with pyruvate showed a rather rapid adaptation, characterized by a steady increase in the stressing/relaxing dilution pulse ratio. On the other hand, the culture fed with glycerol showed a two-phase plateau-acceleration development. We isolated strains from the two cultures on the stressing medium and cultivated them on a minimal medium supplemented with different carbon sources (Fig. 5B). Both strains were able to grow on all carbon sources, with the exception of those entering the TCA cycle – that is succinate and 2-ketoglutarate.

**Figure 5:**
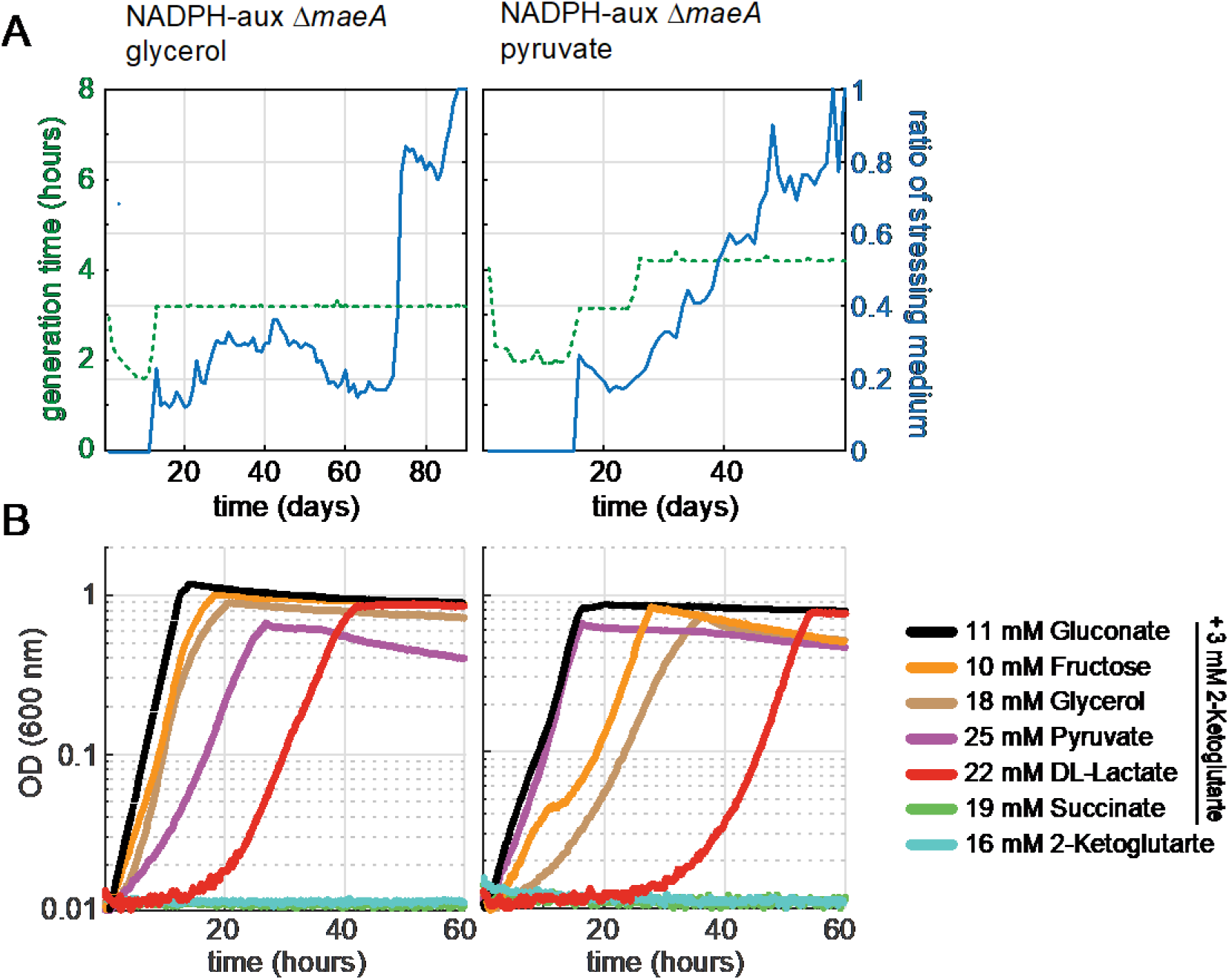
Evolution of the NADPH-auxotroph Δ*maeA* strain for growth on glycerol or pyruvate in the absence of gluconate. For each carbon source 2 independent cultures were subjected to a medium swap regime in GM3 devices (see methods section). For both carbon sources, one of the two cultures evolved to growth in stressing medium. **A** Evolutionary kinetics of the cultures in the GM3 device. The ratio of stressing medium (5 mM 2-ketoglutarate plus glycerol (20mM) or pyruvate (25 mM)) over relaxing medium (same composition as stressing medium plus 5 mM D-gluconate) is shown by the blue line (right axes). Generation times are indicated by the green dashed lines (left axes). **B** Growth of isolated mutants was determined on 11 mM gluconate, 10 mM fructose, 18 mM glycerol, 25 mM pyruvate, 22 mM D/L-lactate, 19 mM succinate (all supplemented with 3 mM 2-ketoglutarate), and 16 mM 2-ketoglutarate. Growth curves were recorded in triplicates, showing similar growth (± 5%).

Genomic sequencing of isolates from the two cultures revealed missense mutations in *lpd* (Table 1 and Supplementary Data). In all isolates, residue E205 was mutated either to glycine or to alanine; in one isolate, the E205A mutation was further accompanied by an E366K mutation. We note that the E205A mutation was also identified following the adaptive evolution of the NADPH-auxotroph strain cultivated on 2-ketoglutarate in which *maeA* did not mutate (see above, Table 1). It therefore seems that Lpd, which participates as a subunit in the pyruvate dehydrogenase and 2-ketoglutarate dehydrogenase complexes [14], was mutated to accept NADP^+^.

We used MAGE to introduce the three observed mutation sets – E205G, E205A, and E205A E366K – to the *lpd* gene in the non-evolved, parental strain (NADPH auxotroph deleted in *maeA*). All resulting strains were found to grow without gluconate on (almost) all carbon sources, where the E205G mutation seems to enable the best growth (Fig. 6). Interestingly, while the isolated strains from the evolved culture could not grow on 2-ketoglutarate and succinate (Fig. 5B), the MAGE-constructed strains could grow on these carbon sources (Fig. 6, with the exception of Lpd E205A cultivated on succinate). This could potentially be attributed to the adaptation of the evolved strains for growth on carbon sources that enter glycolysis and hence force anaplerotic flux (glycerol, pyruvate) rather than enter the TCA cycle.

**Figure 6:**
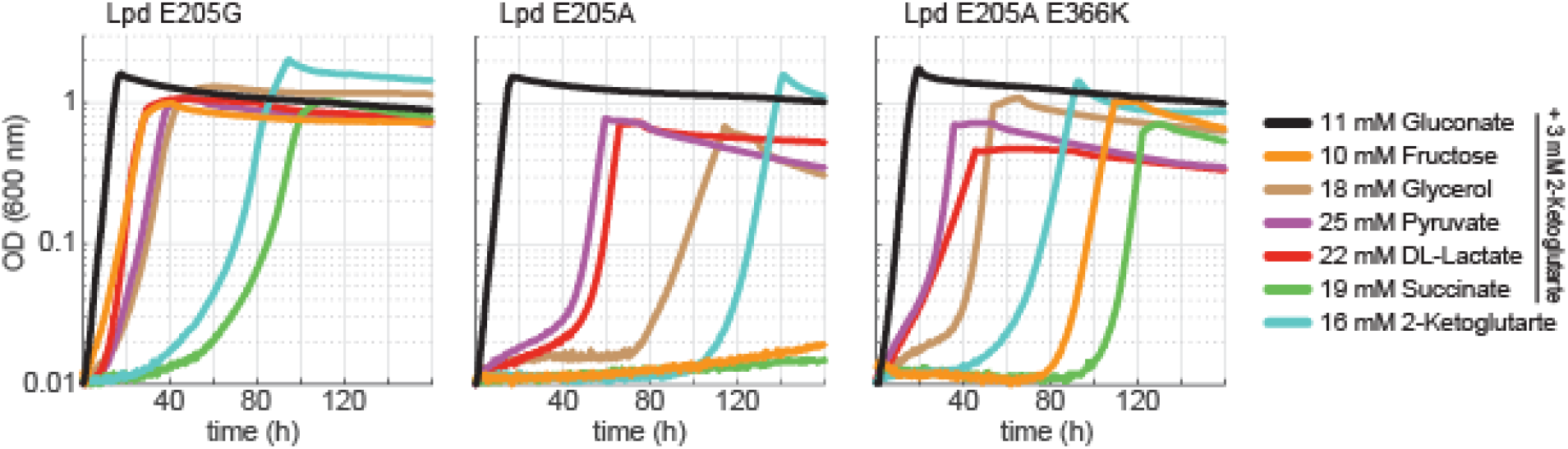
Growth of the NADPH-auxotroph Δ*maeA* strain derivatives mutated in *lpd* (E205G, E205A, E205A + E366K) using the MAGE protocol (see methods section). Growth was determined on 11 mM gluconate, 10 mM fructose, 18 mM glycerol, 25 mM pyruvate, 22 mM D/L-lactate, 19 mM succinate (all supplemented with 3 mM 2-ketoglutarate), and 16 mM 2-ketoglutarate. Curves were recorded in triplicates, showing similar growth (± 5%).

We further characterized the kinetics of the purified Lpd WT, Lpd E205G, Lpd E205A, and Lpd E205G E366K. We found that while Lpd WT displayed no detectable activity with NADP^+^, the mutated Lpd versions catalyzed NADP^+^ reduction with *k*_*cat*_/*K*_*M*_ of 7.8, 6.5, and 10.6 s^-1^ mM^-1^ for Lpd E205G, Lpd E205A, and Lpd E205G E366K, respectively (Table 3). Interestingly, the preference of Lpd E205G, Lpd E205A, and Lpd E205G E366K towards NADP^+^, (*k*_*cat*_/*K*_*M*_)_NADP_/(*k*_*cat*_/*K*_*M*_)_NAD_ between 0.6 and 1.9, is considerably lower than that observed with the mutated variants of MaeA. And yet, such slight preference seems to ensure effective regeneration of NADPH by the mutated Lpds, thus enabling growth in a medium lacking gluconate. This might be explained by the fact that the combined flux via the pyruvate dehydrogenase and 2-ketoglutarate dehydrogenase complexes is substantially higher than that via the malic enzyme, especially since 2-ketoglutarate, as a necessary supplement (due to the *icd* deletion), was present in all experiments.

**Table 3.** Apparent steady state parameters of Lpd variants with NAD^+^ and NADP^+^. Underlying Michaelis-Menten kinetics can be found in Supplementary Figure S2.

| Enzyme | Substrate | $K_M$ (mM) | $K_{cat}$ (s<sup>-1</sup>) | $K_{cat} / K_M$<br>(s<sup>-1</sup> mM<sup>-1</sup>) | $(K_{cat} / K_M)_{NADP} /$<br>$(K_{cat} / K_M)_{NAD}$ |
| --- | --- | --- | --- | --- | --- |
| Lpd WT | dihydrolipoamide | 0.18 ± 0.02 |  |  |  |
|  | NAD<sup>+</sup> | 0.44 ± 0.06 | 363 ± 60 | 830 |  |
|  | NADP<sup>+</sup> | no activity |  |  |  |
| Lpd E205G | dihydrolipoamide | 0.24 ± 0.09 |  |  | 0.6 |
|  | NAD<sup>+</sup> | 9.49 ± 1.82 | 121.2 ± 0.9 | 12.8 |  |
|  | NADP<sup>+</sup> | 3.48 ± 1.03 | 27.2 ± 3.4 | 7.8 |  |
| Lpd E205A | dihydrolipoamide | 0.02 ± 0.00 |  |  | 1.2 |
|  | NAD<sup>+</sup> | 6.35 ± 0.38 | 34.3 ± 0.9 | 5.4 |  |
|  | NADP<sup>+</sup> | 2.48 ± 0.69 | 16.2 ± 2.5 | 6.5 |  |
| Lpd E205A<br>E366K | dihydrolipoamide | 0.03 ± 0.01 |  |  | 1.9 |
|  | NAD<sup>+</sup> | 8.07 ± 0.45 | 45.2 ± 1.3 | 5.6 |  |
|  | NADP<sup>+</sup> | 3.26 ± 0.81 | 34.4 ± 2.9 | 10.6 |  |

Finally, to assess the possibility that other central metabolism oxidoreductases could mutate to enable NADPH regeneration, we deleted both *maeA* and *lpd* genes in the NADPH-auxotroph strain. We subjected the resulting strain (Δ*zwf* Δ*maeB* Δ*icd* Δ*pntAB* Δ*sthA* Δ*maeA* Δ*lpd*) to medium swap adaptation as described above, using either glycerol or succinate as a carbon source (both relaxing and stressing media were further supplemented with acetate to cope with *lpd* deletion). However, even after prolonged cultivation (>1000 generations), none of the four cultures evolved towards growth without gluconate.

## Discussion

In this study we demonstrated whole-cell adaptive evolution towards the emergence of novel routes for NADPH regeneration. In ten successful such experiments, using different carbon sources, we found two NAD-dependent oxidoreductases within central metabolism that have evolved to accept NADP^+^: the malic enzyme MaeA and lipoamide dehydrogenase Lpd.

Notably, the key residue that was mutated in all identified MaeA variants – aspartate 336 (Table 2) – was also found to mutate in a previous study, enabling reductive, CO_2_-assimilating flux using NADPH as an electron donor [15]. The preference towards NADP^+^ of the MaeA D336G variant described before was ≈12, somewhat lower than that observed for our mutants, i.e., 17-18. Moreover, the affinity of MaeA D336G towards NADP^+^, *K*_*M,app*_ = 0.23 mM, was an order of magnitude lower that that observed with our mutants MaeA D336N L176V and MaeA D336A I283N, i.e., *K*_*M,app*_ < 0.02 mM. This emphasizes the supportive role of the L176V and I283N mutations in restoring high catalytic efficiency upon cofactor switching. As the physiological concentration of NADP^+^ is exceedingly low, characteristically ≈ 1 μM in *E. coli* WT [16], sustaining high affinity towards this cofactor is key for its effective reduction. This explains why in most mutated MaeA variants identified, D336 was not mutated alone, but was rather accompanied by another mutation. As mentioned above, the relative catalytic efficiency of MaeA D336N L176V and MaeA D336A I283N with NADP^+^ was found to be 4-fold higher than that of the MaeA WT with NAD^+^, thus representing one of the few cases in which cofactor switching is coupled to improved overall kinetics. An *a priori* identification of the L176 or I283 residues as targets for recovery and increase of catalytic activity would be very difficult, thus demonstrating the power of natural selection in leading to superior, yet non-trivial solutions.

The key residue that was mutated in all three Lpd variants – glutamate 205 (Table 3) – was also previously mutated to switch Lpd cofactor specificity [17]. In this study, the catalytic efficiency with NADP^+^ of the best engineered variant of Lpd, which harbored the E205V mutation and four additional point mutations, was 100-fold higher than that with NAD^+^; and 4-fold superior to the catalytic efficiency of the Lpd WT with NAD^+^.

The consequences of the sole mutation at position 205 (E205V) on the kinetic parameters of the enzyme were not investigated. The selection of the same five combined mutations in an evolutionary experiment in continuous culture would require several events of point mutation to arise and be fixed in the same gene, which makes the occurrence of such a variant highly improbable. Comparatively, the variant Lpd enzymes selected in our experiments exhibited similar catalytic efficiencies with both nicotinamide cofactors (ratio (*k*_*cat*_/*K*_*M*_)_NADP_/(*k*_*cat*_/*K*_*M*_)_NAD_ between 0.6 and 1.9) and far lower catalytic efficiencies with NAD^+^ than Lpd WT. The mutations selected by the process of natural selection enlarged the cofactor specificity of Lpd with a concomitant decrease of activity, which apparently constitutes a good compromise for maintaining balanced pools of NADH and NADPH and for ensuring sufficient carbon flux to sustain growth.

Many of the evolved strains harbor a mutation in the gene coding for citrate synthase (*gltA*, Supplementary Data). Interestingly, we found that even a short-term cultivation of the NADPH-auxotroph strain using a ‘permissive medium’ – having gluconate in the minimal medium – frequently led to mutations in this gene. These mutations included a missense mutation changing residue 150 from leucine to arginine which resulted in a 30-fold decrease in specific activity (Supplementary Table S1) and a base pair deletion causing a frameshift which shortened the polypeptide from 427 AA to 317 AA (Supplementary Data). Citrate synthase activity, while not being required for the NADPH-auxotroph strain in the presence of 2-ketoglutarate, is probably deleterious as it leads to overproduction and accumulation of citrate and isocitrate, which cannot be further metabolized easily due to the deletion of Icd (considering limited flux via the glyoxylate shunt). The downregulation or elimination of GltA activity avoids this overproduction and hence might be beneficial for cell growth.

It is worth noticing that out of all possible oxidoreductase enzymes in central metabolism only MaeA and Lpd were found to mutate during the adaptive evolution. This might be explained by the fact that shifting the cofactor specificity of these two enzymes was attainable thanks to a unique point mutation. Changing the cofactor preference of the other central metabolism oxidoreductases might require the accumulation of multiple mutations, for both cofactor switching and recovery of activity [1, 13]. However, a more plausible option is that the evolution of other oxidoreductases towards NADPH regeneration is biochemically constrained due to unfavorable thermodynamics. For example, the reactions catalyzed by glycerol-3-phosphate dehydrogenase (GpsA) or lactate dehydrogenase (LdhA) strongly favor NADH consumption (Δ_r_G’^m^ < -20 kJ/mol [18]). Hence, even if these enzymes were mutated to accept NADP^+^, their ability to regenerate NADPH with a sufficiently high rate – when cells are fed with the respective carbon source – would be highly limited. Indeed, glycerol and lactate catabolism involve oxidoreductases using a quinone as electron acceptor rather than NAD(P)^+^ (Fig. 1).

Similarly, if glyceraldehyde 3-phosphate dehydrogenase (GapA) would evolve to accept NADP^+^, a severe thermodynamic barrier might arise. Even with NAD^+^, substrate-level phosphorylation (involving GapA, Δ_r_G’^m^ = 24,9 kJ/mol) represents the major thermodynamic barrier in glycolysis [19]. As the cellular NADP pool is substantially more reduced than the NAD pool, switching the cofactor preference of GapA from NAD^+^ to NADP^+^ would further reduce the reaction driving force, rendering glycolysis practically inoperative. Finally, the oxidation of malate to oxaloacetate is the most thermodynamically challenging reaction in central metabolism (Δ_r_G’^m^ = 30,3 kJ/mol), which can operate only if the concentration of oxaloacetate is kept very low (∼1 µM) [19]. Hence, replacing NAD^+^ with NADP^+^, thus further decreasing the driving force for malate oxidation, would certainly render the TCA cycle thermodynamically infeasible. Taken together, it seems that only those central metabolism oxidoreductases which thermodynamically prefer the NAD^+^ reduction direction (MaeA, Δ_r_G’^m^ = -4.1 kJ/mol; Lpd as part of pyruvate dehydrogenase, Δ_r_G’^m^ = -35.3 kJ/mol and as part of 2-ketoglutarate dehydrogenase -27.2 kJ/mol) could evolve to accept NADP^+^, as only these enzymes could sustain a high rate of NADPH regeneration.

Interestingly, instead of evolving a central metabolism oxidoreductase to accept NADP^+^, the adaptive evolution could have increased metabolic flux towards routes that natively produce NADPH but usually carry only low fluxes. For example, increasing flux towards serine and glycine biosynthesis and one carbon metabolism could boost NADPH regeneration via the NADP-dependent bifunctional 5,10-methylene-tetrahydrofolate dehydrogenase/5,10-methenyl-tetrahydrofolate cyclohydrolase (FolD). The fact that we did not observe such adaptation indicates that it is easier to change the cofactor preference of an enzyme via few mutations rather than redistribute fluxes within the endogenous metabolic network.

Overall, the presented work shows the power of evolution and the flexibility, but also the limits of metabolism to adapt to metabolic challenges. Finding a solution counteracting the increased metabolic constraint by the additional deletion of both *maeA* and *lpd* in the NADPH-auxotroph was not within reach in our setup, most likely because of the thermodynamic constraints of the remaining oxidoreductases, which additionally might not provide easy starting points for a cofactor switch to NADP^+^. However, the failure of the evolution attempt of the NADPH-auxotroph Δ*maeA* Δ*lpd* indicates that this strain provides a stringent host for the *in vivo* testing of heterologous NADP^+^ specific oxidoreductases and their evolution.

## Methods

### Reagents and chemicals

Primers were synthesized by Eurofins (Ebersberg, Germany) (Supplementary Table S2). Screening PCRs were performed using DreamTaq polymerase (Thermo Fisher Scientific, Dreieich, Germany). PCR reactions for amplifying deletion cassettes were done using PrimeSTAR MAX DNA Polymerase (Takara). NAD^+^ and NADP^+^(Na)_2_ were purchased from Carl Roth GmbH, malic acid and chicken egg lysozyme from Sigma Aldrich AG and DNAse I from Roche Diagnostics. Dihydrolipoamide was synthesized by borohydride reduction of lipoamide (Sigma Aldrich SA) as described by Reed et al. [20]. Purity (> 95 %) was checked by NMR and infusion mass-spectrometry analysis.

### Media

LB medium (1% NaCl, 0.5% yeast extract, 1% tryptone) was used for strain maintenance. When appropriate, kanamycin (25 μg/mL), ampicillin (100 μg/mL) or chloramphenicol (30 μg/mL) was used. Minimal MA medium (31 mM Na_2_HPO_4_, 25 mM KH_2_PO_4_, 18 mM NH_4_Cl, 1 mM MgSO_4,_ 40 µM trisodic nitrilotriacetic acid, 3 μM CaCl_2_, 3 μM FeCl_3·_6H_2_O, 0.3 μM ZnCl_2_, 0.3 μM CuCl_2_·2H_2_O, 0.3 μM CoCl_2_·2H_2_O, 0.3 μM H_3_BO_3_, 1 μM MnCl_2_, 0.3 µM CrCl_3_, 6 H_2_O, 0.3 µM Ni_2_Cl, 6 H_2_O, 0.3 µM Na_2_MoO_4_, 2 H_2_O, 0.3 µM Na_2_SeO_3_, 5 H_2_O) was used for long-term continuous cultures. M9 minimal medium (50 mM Na_2_HPO_4_, 20 mM KH_2_PO_4_, 1 mM NaCl, 20 mM NH_4_Cl, 2 mM MgSO_4_ and 100 μM CaCl_2_, 134 μM EDTA, 13 μM FeCl_3_·6H_2_O, 6.2 μM ZnCl_2_, 0.76 μM CuCl_2_·2H_2_O, 0.42 μM CoCl_2_·2H_2_O, 1.62 μM H_3_BO_3_, 0.081 μM MnCl_2_·4H_2_O) was used for cell growth analysis. The mineral media were supplemented with various carbon sources as indicated in the main text and hereafter.

### Strains and plasmids

*E. coli* K12 strains used in this study are derivatives of strain SIJ488, which was used as wildtype reference (Table 4). The deletion of the *maeA* gene was carried out by λ-Red recombination using a kanamycin resistance cassette generated via PCR using the FRT-PGK-gb2-neo-FRT (Km) cassette (Gene Bridges, Germany) and the primer pair maeA_KO_fw and maeA_KO_rv. Primers maeA_KO_Ver-F and maeA_KO_Ver-R were used to verify the deletion of *maeA* (Supplementary Table S2). Cell preparation and transformation for gene deletion was carried out as described [21, 22]. The coding sequences of the WT sequences of *maeA, lpd* and *gltA*, as well as the respective mutated genes were amplified by PCR using the primer pairs maeA_Nter_histag_fw and maeA_rv, lpd_Nter_histag_fw and lpd_rv, gltA_Nter_histag_fw and gltA-R, respectively (Supplementary Table S2). The amplified fragments were inserted into a modified Novagen pET22b(+) expression vector (Supplementary Table S3) by using a ligation independent directional cloning method [23]. The sequence of the inserts of the resulting plasmids was verified by Sanger sequencing.

**Table 4:**
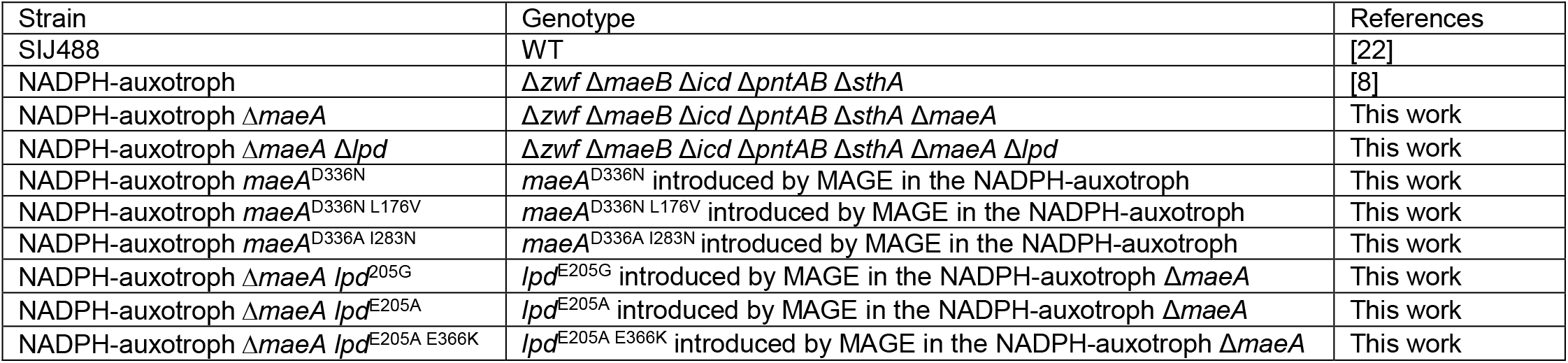
*E. coli* strains and their genetic modifications used in this study.

### Evolution in GM3-driven long-term continuous culture

Pre-cultures of the auxotrophic strains NADPH-auxotroph and NADPH-auxotroph Δ*maeA* were obtained in permissive growth media consisting in minimal MA medium supplemented with 5 mM D-gluconate, 5 mM 2-ketoglutarate and one of the following carbon sources: D-fructose (10 mM), succinate (17 mM), pyruvate (25 mM), lactate (25 mM), glycerol (20 mM), 2-ketoglutarate (20 mM final). Each pre-culture was then used to inoculate the growth chambers (16 ml per chamber) of two parallel independent GM3 devices [10]. A continuous gas flow of sterile air through the culture vessel ensured constant aeration and growth in suspension by counteracting cell sedimentation. The cultures were grown in the corresponding medium under turbidostat mode (dilution threshold set to 80 % transmittance (OD ≈ 0.4, 37°C) until stable growth of the bacterial population. The cultures were then submitted to conditional medium swap regime. This regime enabled gradual adaptation of the bacterial populations to grow in a non-permissive or stressing medium of composition equivalent to the permissive medium but lacking D-gluconate. Dilutions of the growing cultures were triggered every 10 minutes with a fixed volume of medium calculated to impose a generation time of 3h10 on the cell population, if not otherwise stated. The growing cultures were fed by permissive or stressing medium depending on the turbidity of the culture with respect to a set OD threshold (OD_600_ value of 0.4). When the OD exceeded the threshold, a pulse of stressing medium was injected; otherwise a pulse of permissive medium. The cultures were maintained under medium swap regime until the bacterial cell populations grew in 100 % stressing medium. Cultures which did not evolve towards growth in stressing medium were aborted after culturing for 1000 generations. Four isolates were obtained on agar-solidified stressing medium for each successful evolution experiment and further analyzed.

### Genomic analysis of evolved strains

Pair-end libraries (2×150 bp) were prepared with 1 µg genomic DNA from the evolved strains and sequenced using a MiSeq sequencer (Illumina). The PALOMA pipeline, integrated in the platform Microscope (http://www.genoscope.cns.fr/agc/microscope) was used to map the reads against *E. coli* K12 wildtype strain MG1655 reference sequence (NC_000913.3) for detecting single nucleotide variations, short insertions or deletions (in/dels) as well as read coverage variations [24].

### Growth experiments

Overnight cultures were obtained in 4 mL M9 medium supplemented with 12 mM gluconate and 3 mM 2-ketoglutarate (permissive growth condition). Strains were harvested (6,000*g*, 3 min) and washed thrice in M9 medium to remove residual carbon sources. Cells were then inoculated into the various test media to OD_600_ of 0.01 and distributed into 96-well microtiter plates (Nunclon Delta Surface, Thermo Scientific). Each well contained 150 μL of culture and 50 μL mineral oil (Sigma-Aldrich) to avoid evaporation. Growth monitoring and incubation at 37 °C was carried out in a microplate reader (EPOCH 2, BioTek). The program (controlled by Gen5 3.04) consisted in 4 shaking phases, 60 seconds each: linear shaking 567 cpm (3 mm), orbital shaking 282 cpm (3 mm), linear shaking 731 cpm (2 mm), orbital shaking 365 cpm (2 mm). After 3 shaking cycles absorbance OD_600_ was measured. Raw data were calibrated to 1 cm-wide cuvette measured OD_600_ values according to OD_cuvette_ = OD_plate_ / 0.23. Matlab was used to calculate growth parameters, repeatedly based on at least three technical replicates. Average values were used to generate the growth curves. Variability between triplicate measurements was less than 5% in all cases displayed.

### Reverse engineering

The pORTMAGE system which allows an efficient directed genome editing in *E. coli* [25] was used to introduce into the naïve ancestor strains the mutations fixed in the genes *maeA* and *lpd* during the evolution experiments. MAGE oligos were designed using http://modest.biosustain.dtu.dk/ (Supplementary Table S2); they contained thioester bonds at 5’ and 3’ ends and the wanted mutation. Cells carrying the pORTMAGE-2 plasmid were incubated at 30°C. When cultures reached an OD_600_ of 0.5, the system was induced by incubation at 42°C for 15 min. Afterwards cells were immediately chilled on ice until they were prepared for electroporation by 3 consecutive cycles of washing and centrifugation (11,000 rpm, 30 sec, 2°C) with ice-cold 10 % glycerol solution. MAGE oligos were introduced into the strains by electroporation (1 mm cuvette, 1.8 kV, 25 µF, 200 Ω). Strains were directly transferred to LB, 10 mM gluconate, 3 mM 2-ketoglutarate and incubated for 1 hour. After three rounds of MAGE cells were plated on LB-plates containing 10 mM gluconate and 3 mM 2-ketoglutarate. The respective loci were amplified by PCR using respective primer pairs Ver-F and Ver-R (Supplementary Table S2), and sequenced by Sanger sequencing, to identify strains with the wanted mutations.

### Protein Expression and purification

The His-tagged WT and mutated MaeA proteins were expressed in *E. coli* BL21 DE3. Cells in Terrific Broth containing 100 µg/mL ampicillin were grown at 37 °C until they reached an OD_600_ of 0.8-1 upon which expression for 16 h at 23 °C was induced by addition of 250 μM IPTG (Isopropyl-D-β-thiogalactopyranoside). Cells were harvested for 15 min at 6’000 g at 4°C then resuspended in 2 mL of Buffer A (50 mM Tris, 500 mM NaCl, pH 7.5) per gram of pellet. The suspension was treated with 10 mg/mL of DNAse I, 5 mM MgCl_2_ and 6 µg/mL lysozyme on ice for 20 min upon which cells were lysed by sonication. The lysate was clarified at 45’000 g at 4°C for 45 min and the supernatant was filtered through a 0.4 µm syringe tip filter (Sarstedt, Nümbrecht, Germany). Lysate was loaded onto a pre-equilibrated 1 mL HisTrap FF column and washed with 12 % Buffer B (50 mM Tris, 500 mM NaCl, 500 mM imidazole, pH 7.5) for 20-30 column volumes until the UV 280 nm reached the baseline level. The protein was eluted by applying 100% buffer B, collected then pooled and desalted into 12.5 mM Tris, 125 mM NaCl, pH 7.5, 10 % glycerol. The protein was frozen in N_2_ (l) and stored at -80°C if not immediately used for assays.

The His-tagged WT and mutated Lpd proteins were expressed in *E. coli* BL21 DE3 Codon+ (Invitrogen). Cells in 400 ml Terrific broth containing 100 µg/mL carbenicillin were grown at 37 °C until they reached an OD_600_ = of 0.8-1 upon which expression for 16 h at 20 °C was induced by addition 500 μM IPTG. Cells were harvested by centrifugation for 30 min at 10000 g at 4°C. Cell pellets were frozen at -80°C for one night. Thawed cells were then suspended in 32 ml of Buffer A (50 mM phosphate (Na/K), 500 mM NaCl, 30 mM imidazole, 15% glycerol, pH 8.0) and lysed for 30 min at room temperature after addition of 3.6 ml of Bug Buster (Novagen) 32 µl DTT (dithiothreitol) 1M, 320 µl Pefabloc 0.1 M (Millipore) and 23 µl Lysonase (Novagen). Lysate was clarified at 9000g for 45 min at 4°C then loaded onto a 5 ml HisTrap FF column pre-equilibrated in Buffer A. The protein was eluted in Buffer B (50 mM phosphate (Na/K), 500 mM NaCl, 250 mM imidazole, 1 mM DTT 15% glycerol, pH 8.0) and desalted on a gel-filtration column Hi Load 16/60 Superdex 200 pg in Buffer C (50mM Tris, 50 mM NaCl, glycerol 15%, 1 mM DTT, pH8.0). The protein was frozen and stored at -80°C if not immediately used for assays.

### Biochemical assays

#### Characterization of MaeA kinetic parameters

Assays were performed on a Cary-60 UV/Vis spectrophotometer (Agilent) at 30°C using quartz cuvettes (10 mm path length; Hellma). Reactions were performed in 50 mM Tris HCl pH 7.5 10 mM MgCl_2_. Kinetic parameters for one substrate were determined by varying its concentration while the others were kept constant at 6-10 times their K_M_ value. Reaction procedure was monitored by following the reduction of NAD(P)^+^ at 340 nm (ε_NADPH,340nm_ = 6.22 mM^-1^ cm^-1^). Each concentration was measured in triplicates and the obtained curves were fit using GraphPad Prism 8. Hyperbolic curves were fit to the Michaelis-Menten equation to obtain apparent k_cat_ and K_M_ values.

#### Characterization of Lpd kinetic parameters

Assays were performed using a Safas UV mc2 double beam spectrophotometer at room temperature using quartz cuvettes (10mm path length). The concentration of purified Lpd enzyme was determined spectrophotometrically using an extinction coefficient of 34.0 mM^- 1^.cm^-1^ at 280 nm. The concentration of FAD was determined using an extinction coefficient of 15.4 mM^- 1^.cm^-1^ at 446 nm [26]. The absorbances at 446 nm and at 280 nm were measured and the ratio calculated to determine the fraction of active FAD-containing catalysts within each batch of purified enzyme and to normalize the results between the different enzyme forms. Assays of Lpd-catalyzed oxidation of dihydrolipoamide were conducted in 100 mM Na phosphate, 100 mM KCl, 8 mM TCEP pH 7.6. Kinetic parameters for NAD(P)+ were determined by varying its concentration in the presence of a saturating concentration of dihydrolipoamide (4 mM). The reactions were monitored by recording the accumulation of NAD(P)H at 340 nm. Kinetic constants were determined by non-linear analysis of initial rates from duplicate experiments using SigmaPlot 9.0 (Systat Software, Inc.).

## Data availability

The data supporting the findings of this work are available within the paper and its Supplementary Information files. Strains used here are available on request from the corresponding author. For ΔG calculations data from eQuilibrator (http://equilibrator.weizmann.ac.il/) was used.

## Acknowledgements

We express our gratitude to the late Arren Bar-Even for his guidance and unwavering support. This study was funded by the Max Planck Society and the CEA Genoscope.

## Author contributions

S.N.L. and A.B.-E. conceived the study. M.B., V.D., A.P., T.E., S.N.L., and A.B.-E. designed the experiments. M.B. and V.D. supervised the evolution experiments and analyzed the genome sequencing data. I.D. and A.B. ran the continuous cultures, isolated and characterized evolved strains. D.R. performed comparative genomic analysis. S.N.L., S.M., L.C.R. performed reverse engineering and growth experiments. G.S., L.C.R., M.F., A.B., A.P. and T.E. performed the *in vitro* experiments. S.N.L., A.B.-E, V.D., and M.B. analyzed the results and wrote the manuscript with contributions from all authors.

## Supplementary material

Figure S1: Michaelis-Menten kinetics of MaeA variants. Data represents mean values +/- SD from three independent experiments (n = 3).

Figure S2: Michaelis-Menten kinetics of LPD variants. Data represents mean values +/- SD from three independent experiments (n = 3).

Table S1: Citrate synthase activity.

Table S2: List of DNA oligo primers used in this study. *thioester bound

Table S3. Plasmids constructed for protein purification

## Supplementary Methods

### GltA expression and purification

The His-tagged WT and L150R-mutated GltA proteins were expressed in *E. coli* BL21 DE3 Codon+ (Invitrogen). Cells in Terrific broth containing 100 µg/mL carbenicillin were grown at 37 °C until they reached an OD_600_ = of 0.8-1 upon which expression for 16 h at 20 °C was induced by addition of 500 μM IPTG. Cells were harvested by centrifugation for 30 min at 10000g at 4°C. Cell pellets were frozen at -80°C for one night. Thawed cells were then suspended in 3 ml of Buffer A (50 mM HEPES, 50 mM NaCl, 30 mM imidazole, 10% glycérol, pH 8.0) and incubated with Pefabloc and Lysonase for 20 min then lysed by sonication. The lysate was clarified at 12000 g for 30 min at 4 °C. The supernatant was loaded onto a pre-equilibrated Ni-NTA minicolumn (QIAGEN) and washed thrice with Buffer A. The protein was eluted in elution buffer B (50 mM HEPES, 50 mM NaCl 250 mM imidazole, 10 % glycerol), collected, pooled and desalted on a Amicon Ultra-4 10kD column in buffer C (50 mM HEPES 50 mM NaCl 10 % glycerol). The protein was frozen and stored at -80°C if not immediately used for assays.

### Measurement of GltA specific activity

Citrate synthase activity was determined by measuring the initial rate of reaction at 412 nm by means of the DTNB method [1]. Reactions were conducted in 100 mM Tris pH 8, 200 µM DTNB, sub-saturating concentrations of acetyl-CoA (200 µM) and oxaloacetate (100 µM) and with or without the addition of NADH (200 µM) and KCl (100 mM). Specific activity (µmole min^-1^ mg^-1^) was determined.

## References

1. Cahn, J. K., Werlang, C. A., Baumschlager, A., Brinkmann-Chen, S., Mayo, S. L. & Arnold, F. H. (2017) A General Tool for Engineering the NAD/NADP Cofactor Preference of Oxidoreductases, ACS synthetic biology. 6, 326–333.

2. Chanique, A. M. & Parra, L. P. (2018) Protein Engineering for Nicotinamide Coenzyme Specificity in Oxidoreductases: Attempts and Challenges, Frontiers in microbiology. 9, 194.

3. Selles Vidal, L., Kelly, C. L., Mordaka, P. M. & Heap, J. T. (2018) Review of NAD(P)H-dependent oxidoreductases: Properties, engineering and application, Biochimica et biophysica acta Proteins and proteomics. 1866, 327–347.

4. Wang, M., Chen, B., Fang, Y. & Tan, T. (2017) Cofactor engineering for more efficient production of chemicals and biofuels, Biotechnology advances. 35, 1032–1039.

5. Zhang, L., King, E., Luo, R. & Li, H. (2018) Development of a High-Throughput, In Vivo Selection Platform for NADPH-Dependent Reactions Based on Redox Balance Principles, ACS synthetic biology. 7, 1715–1721.

6. Kramer, L., Le, X., Rodriguez, M., Wilson, M. A., Guo, J. & Niu, W. (2020) Engineering Carboxylic Acid Reductase (CAR) through a Whole-Cell Growth-Coupled NADPH Recycling Strategy, ACS synthetic biology. 9, 1632–1637.

7. Calzadíaz Ramirez, L., Calvo Tusell, C., Stoffel, G. M., Lindner, S. N., Osuna, S., Erb, T. J., Garcia-Borràs, M., Bar-Even, A. & Acevedo-Rocha, C. G. (2020) In vivo selection for formate dehydrogenases with high efficiency and specificity towards NADP+, ACS Catal. 10, 7512–7525.

8. Lindner, S. N., Ramirez, L. C., Krusemann, J. L., Yishai, O., Belkhelfa, S., He, H., Bouzon, M., Doring, V. & Bar-Even, A. (2018) NADPH-Auxotrophic E. coli: A Sensor Strain for Testing in Vivo Regeneration of NADPH, ACS synthetic biology. 7, 2742–2749.

9. Sauer, U., Canonaco, F., Heri, S., Perrenoud, A. & Fischer, E. (2004) The soluble and membrane-bound transhydrogenases UdhA and PntAB have divergent functions in NADPH metabolism of Escherichia coli, J Biol Chem. 279, 6613–9.

10. Marliere, P., Patrouix, J., Doring, V., Herdewijn, P., Tricot, S., Cruveiller, S., Bouzon, M. & Mutzel, R. (2011) Chemical evolution of a bacterium’s genome, Angew Chem Int Ed Engl. 50, 7109–14.

11. Bouzon, M., Perret, A., Loreau, O., Delmas, V., Perchat, N., Weissenbach, J., Taran, F. & Marliere, P. (2017) A Synthetic Alternative to Canonical One-Carbon Metabolism, ACS synthetic biology. 6, 1520–1533.

12. Wang, H. H., Isaacs, F. J., Carr, P. A., Sun, Z. Z., Xu, G., Forest, C. R. & Church, G. M. (2009) Programming cells by multiplex genome engineering and accelerated evolution, Nature. 460, 894–8.

13. Tawfik, D. S. (2014) Accuracy-rate tradeoffs: how do enzymes meet demands of selectivity and catalytic efficiency?, Curr Opin Chem Biol. 21, 73–80.

14. Carothers, D. J., Pons, G. & Patel, M. S. (1989) Dihydrolipoamide dehydrogenase: functional similarities and divergent evolution of the pyridine nucleotide-disulfide oxidoreductases, Arch Biochem Biophys. 268, 409–25.

15. Zelle, R. M., Harrison, J. C., Pronk, J. T. & van Maris, A. J. (2011) Anaplerotic role for cytosolic malic enzyme in engineered *Saccharomyces cerevisiae* strains, Appl Environ Microbiol. 77, 732–8.

16. Bennett, B. D., Kimball, E. H., Gao, M., Osterhout, R., Van Dien, S. J. & Rabinowitz, J. D. (2009) Absolute metabolite concentrations and implied enzyme active site occupancy in Escherichia coli, Nat Chem Biol. 5, 593–9.

17. Bocanegra, J. A., Scrutton, N. S. & Perham, R. N. (1993) Creation of an NADP-dependent pyruvate dehydrogenase multienzyme complex by protein engineering, Biochemistry. 32, 2737–40.

18. Flamholz, A., Noor, E., Bar-Even, A. & Milo, R. (2012) eQuilibrator--the biochemical thermodynamics calculator, Nucleic Acids Res. 40, D770–5.

19. Noor, E., Bar-Even, A., Flamholz, A., Reznik, E., Liebermeister, W. & Milo, R. (2014) Pathway thermodynamics highlights kinetic obstacles in central metabolism, PLoS Comput Biol. 10, e1003483.

20. Reed, L. J., Koike, M., Levitch, M. E. & Leach, F. R. (1958) Studies on the nature and reactions of protein-bound lipoic acid, J Biol Chem. 232, 143–58.

21. Wenk, S., Yishai, O., Lindner, S. N. & Bar-Even, A. (2018) An Engineering Approach for Rewiring Microbial Metabolism, Methods Enzymol. 608, 329–367.

22. Jensen, S. I., Lennen, R. M., Herrgard, M. J. & Nielsen, A. T. (2015) Seven gene deletions in seven days: Fast generation of Escherichia coli strains tolerant to acetate and osmotic stress, Scientific reports. 5, 17874.

23. Aslanidis, C. & de Jong, P. J. (1990) Ligation-independent cloning of PCR products (LIC-PCR), Nucleic Acids Res. 18, 6069–74.

24. Vallenet, D., Calteau, A., Dubois, M., Amours, P., Bazin, A., Beuvin, M., Burlot, L., Bussell, X., Fouteau, S., Gautreau, G., Lajus, A., Langlois, J., Planel, R., Roche, D., Rollin, J., Rouy, Z., Sabatet, V. & Medigue, C. (2020) MicroScope: an integrated platform for the annotation and exploration of microbial gene functions through genomic, pangenomic and metabolic comparative analysis, Nucleic Acids Res. 48, D579–D589.

25. Nyerges, Á., Csörgő, B., Nagy, I., Bálint, B., Bihari, P., Lázár, V., Apjok, G., Umenhoffer, K., Bogos, B., Pósfai, G. & Pál, C. (2016) A highly precise and portable genome engineering method allows comparison of mutational effects across bacterial species, Proceedings of the National Academy of Sciences. 113, 2502–2507.

26. Thorpe, C., Matthews, R. G. & Williams, C. H., Jr. (1979) Acyl-coenzyme A dehydrogenase from pig kidney. Purification and properties, Biochemistry. 18, 331–7.

## Supplementary reference

1. Moriyama, T. & Srere, P. A. (1971) Purification of rat heart and rat liver citrate synthases. Physical, kinetic, and immunological studies, J Biol Chem. 246, 3217–23.

